# Experimental evolution and genome data analysis of *Candida albicans* reveals cryptic bacteria in single yeast colonies

**DOI:** 10.1101/168500

**Authors:** Danielle do Carmo Ferreira Bruno, Thais Fernanda Bartelli, Camila Ronqui Rodrigues, Marcelo R. S. Briones

## Abstract

At least 25% of patients with positive *Candida albicans* bloodstream infection also have one or more bacterial species associated with the infection. These polymicrobial infections are usually caused by coagulase-negative staphylococci, most commonly *Staphylococcus epidermidis* and are associated with significantly worse clinical outcomes as compared to monomicrobial infections. Here we show bacteria are present in *C. albicans* cultures started from isolated single colony platting. These co-evolving bacteria can only be detected by the use of specific selective medium and/or long periods of incubation from 8 days up to 48 weeks (approximately 4,000 generations), used in experimental evolution methods. The detection of these co-evolving bacteria is highly dependent on the type of enzyme used for 16S rRNA gene amplification and is often missed in clinical laboratory analysis because of short incubation periods, media and temperatures, used in mycology clinical routine, that are unfavorable for bacterial growth. In this study, we identified bacteria in cultures of different *C. albicans* isolates from long term, continuous growth by molecular analysis and microscopy. Also, we confirmed the presence of these co-evolving bacteria by identification of *S. epidermidis* genome segments in sequencing reads of the *C. albicans* reference strain SC5314 genome sequencing project raw data deposited in GenBank. This result rules out the possibility of laboratory specific contamination. Also, we show that the presence of associated bacteria correlates with antifungal resistance alterations observed in growth under hypoxia. Our findings show the intense interaction between *C. albicans* yeasts and bacteria and have direct implications in yeast clinical procedures, especially concerning patient treatment.

## INTRODUCTION

The human commensal fungus *Candida albicans* is part of the microbiota in healthy individuals and colonize several niches, such as skin, gastrointestinal and urogentinal tracts. In immunosuppressed hosts, however, *C. albicans* causes opportunistic mucosal and, often fatal, bloodstream infections (Brown et al., 2012). *C. albicans* coexists with a highly diverse human bacterial microbiota, although, as compared to bacteria, fungi is a small part of the microbiota (Arumugam et al., 2011). These communities often produce mixed species biofilms which have a significant impact in the survival and reproductive success of both bacteria and fungi. These polymicrobial communities are involved in either antagonistic or synergistic relationships (Ovchinnikova et al., 2012). Although most infections by commensal microorganisms originate from endogenous colonization, exogenous contamination also occur, such as infections derived from hospital staff, biofilm contaminated invasive devices or the hospital environment itself (Douglas, 2002; Sims et al., 2005; Vilanova and Correia, 2008). Candidemias are often accompanied by bacterial infections (25%) and this number is believed to be significantly underestimated due to technical difficulties in cultivating bacteria and fungi from a single blood sample (Klotz et al., 2007). Bacteria such as *P. aeruginosa* and the genus *Staphylococcus*, especially *S. epidermidis* and *S. aureus*, are commonly isolated from infections with *C. albicans*. These pathogens are able to colonize the host or medical devices together, forming mixed biofilms that are associated with increased antimicrobial resistance and virulence (Pierce, 2005; Schlecht et al., 2015). However, despite the high incidence and severity, there are still few studies on polymicrobial infections (Schlecht et al., 2015).

To effectively colonize and infect its hosts, *C. albicans* has to adapt to several constrains, including physical barriers such as oxygen levels and temperature, and biological conditions such as carbon sources, nutrient availability, other microbial species and the immune system. Usually in laboratory culturing, micro-environmental conditions that favor microorganismal growth are emphasized, instead of trying to mimic conditions found in the host. While atmospheric oxygen tension, normoxia, is around 21%, in the human body the conditions are hypoxic (between 2 and 9% O_2_) and depending on anatomical location and inflammation the oxygen levels can be as low as ≤1% (Grahl et al., 2012; Setiadi et al., 2006). Temperature is also one of the major barriers to most fungal species that infect mammals (Bergman and Casadevall, 2010). Human commensals, such as *C. albicans*, which are adapted to temperatures near 37°C in the human body, have a cellular adaptation response to temperature increase, which may favor, for example, their survival in infected patients with fever. Therefore, the response to heat shock and hypoxia have a direct impact in adaptation and promotion of infections, especially in the case of nosocomial infections in which the exogenous fungus, living at 21% O_2_ and approximately 23°C, is suddenly transferred to 2-9% O_2_ and 37°C. Another major adaptive advantage of *C. albicans* is its metabolic flexibility. The amount and type of nutrients available vary widely in the human body, depending on the anatomical site (Brock, 2009; Mayer et al., 2013; Rodaki et al., 2009). Once in the bloodstream, *C. albicans* can colonize any organ, where glucose concentration may become limited, although it is assumed that tissue destruction during infection leads to diffusion of new substrates and creation of its own microenvironment (Brock, 2009).

Most studies evaluating antifungal resistance, for example, do not take into account host characteristics, such as temperature, oxygen stresses, other microorganisms in the culture, and nutrient sources. Therefore, evaluating virulence mechanisms in conditions that mimic the host might have an important impact on clinical treatment studies. It is assumed that an agar growing yeast colony is formed exclusively by *Candida spp.* clones (Pujol et al., 1993). Here we show that cultures derived from single isolated colonies of *C. albicans* grown in proper media and conditions reveal the presence of bacteria. This result is confirmed by the analysis of *C. albicans* genomic reads (SC5314 strain), published and deposited in GenBank by other labs (Muzzey et al., 2013), which clearly contain bacterial reads of the same species found in isolates here analyzed. We also show that conditions that mimic the host, such as hypoxia and polymicrobial cultures affect, *in vitro*, the drug susceptibility patterns of *C. albicans*.

## RESULTS

### Identification of co-isolated bacteria in colonies of *C. albicans.*

We were able to identify bacterial DNA in samples from single colonies of *C. albicans*, strain SC5314, in plates were no visual sign of bacterial contamination was observed (Figure 1). We found that media frequently used for yeast growth, such as YPD and *CHROMagar* and culturing for 48h does not reveal bacterial growth, either by colony plating (Figure 2), visualizing aliquots of liquid cultures by light microscopy or by bacterial 16S rRNA gene amplification (Figure 3). Also, cultures grown over a prolonged period (8 days) in YPD liquid medium did not show bacteria when visualized by microscopy, but bacterial DNA could be detected by 16S rRNA gene amplification. Bacteria were detected only under conditions that favor bacterial growth, such as LB, and blood agar, or non-fermentative media such as YPG, at 37°C for 48h (Figure 3). In cultures where bacteria had not been identified by plating or 16S rRNA gene amplification, bacteria were detected by Gram staining microscopy. Gram staining revealed bacterial cells adhered, especially to *C. albicans* hyphae, both in cultures grown for 48h and in 8 days, in YPD, LB, blood agar and *CHROmagar* (Figure 2).

**Figure 1.**
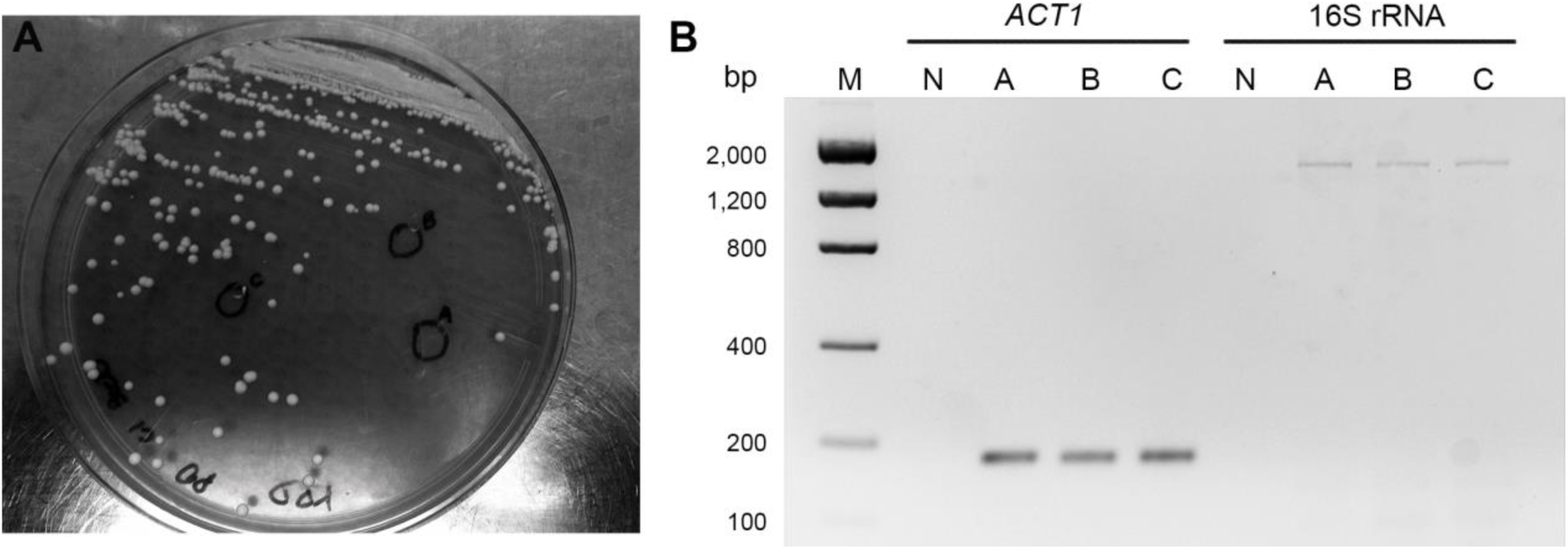
Amplification of *C. albicans ACT1* and bacterial 16S rRNA from a single yeast colony. **A:** *C. albicans* SC5314 YPD plate grown for 48h at 28°C. **B**: *C. albicans ACT1* and bacterial 16S rRNA gene amplification from the 3 single yeasts colonies (*A*, *B* and C) circled in panel **A**. Colonies were inoculated into LB medium and incubated for 8 days at 37°C.

**Figure 2.**
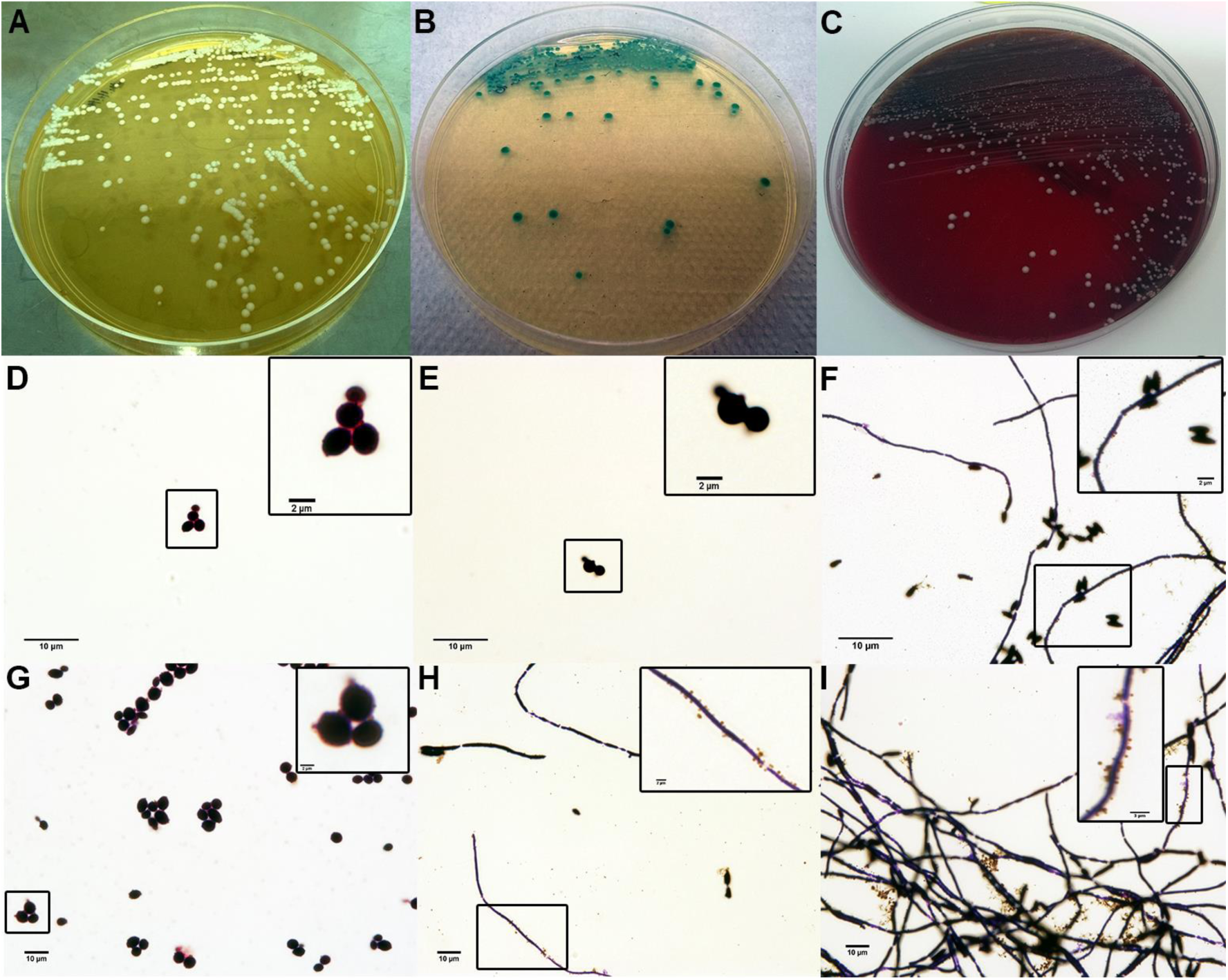
Bacteria present in cultures of *C. albicans* from an isolated colony. Cells of SC5314 grown for 48h and 8 days in different culture media. A: YPD at 28°C, B: *CHROmagar* at 37°C, C: agar blood at 37°C for 48h. D-F: cells grown for 48h and G-I: cells grown for 8 days, in liquid media (YPD for YPD and LB for plates of blood agar and *CHROmagar*) from one colony isolated from the respective plates (A, B and C). Cells were Gram stained and observed in 100x objective with immersion oil.

**Figure 3.**
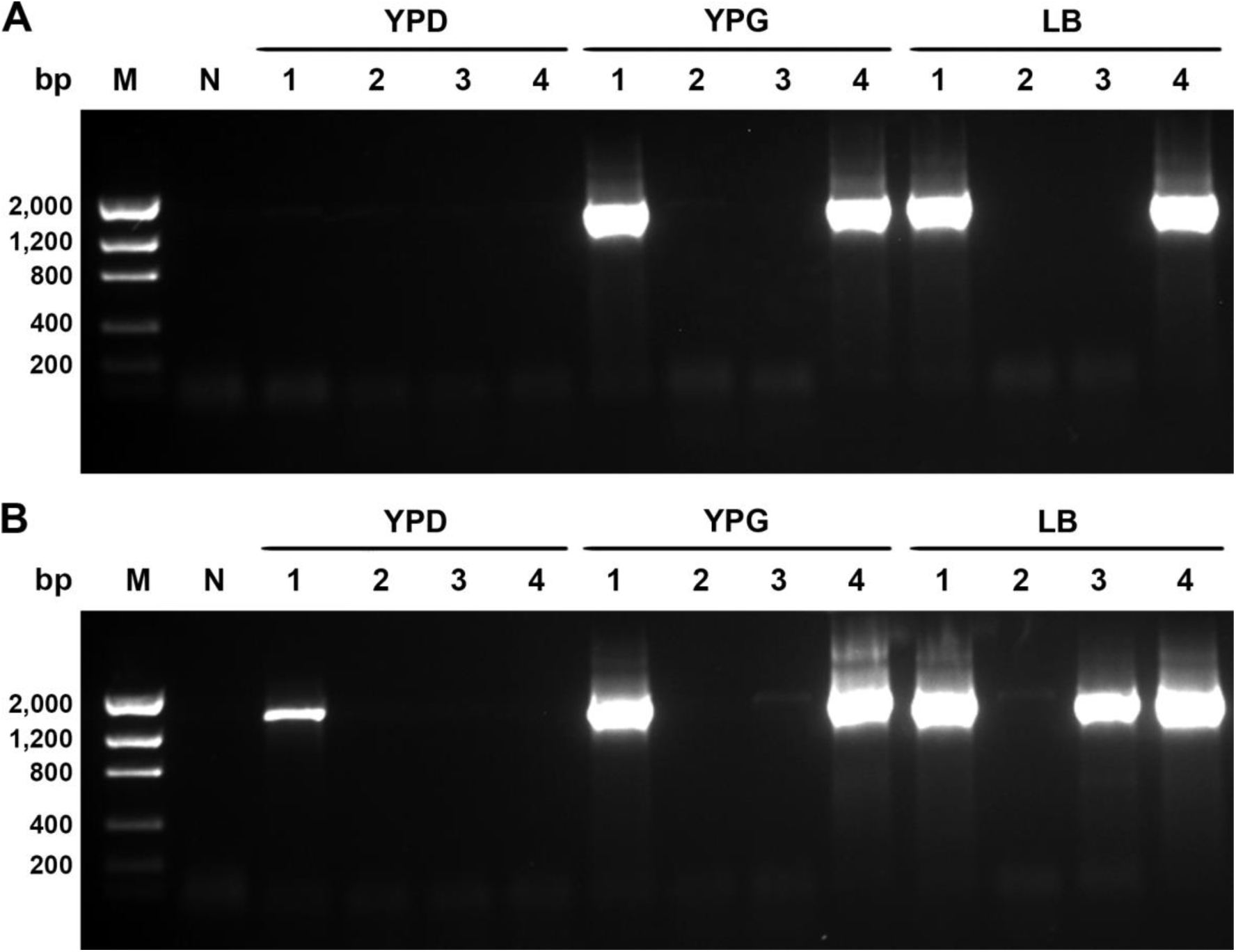
Bacterial 16S rRNA gene amplification in cultures of different *C. albicans* strains in different media. **A**: cultures grown in YPD, YPG and LB for 48h. **B**: cultures grown for 8 days. **M**: Low mass ladder. **N**: negative control. Numbers represent strains. **1**: SC5314. **2**: L757. **3**: ATCC 24433. **4**: ATCC 90029. Amplicons were visualized on 1% w/v agarose gel stained with ethidium bromide.

In addition to strain SC5314, four different *C. albicans* strains (Table 1) were inoculated in YPD, YPG and LB media for 48h or 8 days at 28°C (YPD) and 37°C (other media) (Figure 3). Among the five different *C. albicans* strains tested, L757 was the only that did not show sufficient 16S rRNA gene amplification after 8 days (Figure 3). However, after 12 weeks of continuous growth in YPG medium at 37°C under hypoxic condition, named GTH (Bartelli et al., 2017), it was possible to amplify 16S rRNA for sequencing and species identification. Sequencing of the amplified 16S rRNA gene from total DNA extracted from cultures and BLASTn searches in the 16S ribosomal RNA sequences database, revealed different bacterial species (Table 1). Strains L757 and ATCC 24433 showed the presence of *Propionibacterium acnes* (e-value 0, query coverage 100, identity 99). On the other hand, 16S rRNA sequencing from SC5314, 9117 and ATCC 90029 strains revealed different bacteria of genus *Staphylococcus* (e-value 0, query coverage >99% and identity >97%). Species identification within genus *Staphylococcus* was obtained by sequencing the *Tuf* gene (Heikens et al., 2005). The *Tuf* sequence analysis in strain SC5314 revealed that *S. epidermidis* was present (e-value 0, query coverage 100, identity 100).

**Table 1.** *C. albicans* strains and bacteria species co-isolated as identified by 16S rRNA gene sequencing. ^∗^ Species confirmed by additional sequencing of the gene *Tuf*.

| ***C. albicans***<br>**strains** | **Isolation**<br>**site** | **Geographic**<br>**site** | **Year** | **Bacterial co-isolated** |
| --- | --- | --- | --- | --- |
| **SC5314** | Blood | EUA | 1984 (Gillum et al., 1984) | *Staphylococcus epidermidis* * |
| **L757** | Blood | Brazil | 2001 | *Propionibacterium acnes* |
| **9117** | Blood | Brazil | 2012 | *Staphylococcus epidermidis* |
| **ATCC**<br>**24433** | Nail | EUA | 1994 (Pfaller et al., 1994) | *Propionibacterium acnes* |
| **ATCC**<br>**90029** | Blood | EUA | 1992 (Espinel-Ingroff et al., 1992) | *Staphylococcus warneri* |

Although the results of 16S rRNA sequencing have not suggested more than one genus of bacteria in *C. albicans* strains, we indentified two different bacterial morphologies, coccus and bacillus by electron microscopy (SEM) (Figure 5).

To test whether the published *C. albicans* reference genome (strain SC5314) might contain bacterial genome segments we examined the raw sequencing data available for the *C. albicans* SC5314 genome generated by Muzzey and collaborators (Muzzey et al., 2013) available as SRP022363 NIH SRA. According to these authors, prior to sequencing, strain SC5314 and 9 other homozygous *C. albicans* strains were grown overnight in YPD medium from single colonies and 36 nucleotide paired-end reads sequenced on an Illumina Genome Analyzer IIx, generating 10 runs with approximately 40 million reads each. Sequencing reads were mapped simultaneously with the fasta sequence available for *C. albicans* SC5314 (assembly 22) and the genome sequence available for *S. epidermidis* ATCC 12228 (Zhang et al., 2003), ignoring and excluding reads with non-specific matches. Interestingly, we identified numerous bacterial reads in the raw sequence data that mapped uniquely to the chromosome and plasmid sequences of *S. epidermidis* reference sequence (Tables 2 and 3).

**Table 2.** Mapping reads from raw sequencing data published by Muzzey et al. 2013 (Muzzey et al., 2013) to *C. albicans* (assembly 22, SC5314) and *S. epidermidis* ATCC 12228 (Zhang et al., 2003) reference sequences. Reads with non-specific matches were ignored and not included in the final mapping. The different plasmid samples of pSE-12228 correspond to the different GenBank sequences NC005008.1, NC005007.1, NC005006.1, NC005005.1, NC005004.1, NC005003.1 from top to bottom respectively.

| Reference Sequence | Reference length (bp) | Consensus length (bp) | Fraction of reference covered | Mapped reads | Forward reads | Reversed reads |
| --- | --- | --- | --- | --- | --- | --- |
| *S. epidermidis* chromosome | 2499279 | 1309998 | 0.52 | 275312 | 138004 | 137308 |
| *S. epidermidis* plasmid pSE-12228-01 | 4439 | 2776 | 0.62 | 950 | 418 | 532 |
| *S. epidermidis* plasmid pSE-12228-02 | 4679 | 2693 | 0.57 | 497 | 246 | 251 |
| *S. epidermidis* plasmid pSE-12228-02 | 8007 | 4397 | 0.55 | 562 | 282 | 280 |
| *S. epidermidis* plasmid pSE-12228-02 | 17261 | 10448 | 0.60 | 2887 | 1459 | 1428 |
| *S. epidermidis* plasmid pSE-12228-02 | 24365 | 14221 | 0.58 | 2938 | 1536 | 1402 |
| *S. epidermidis* plasmid pSE-12228-02 | 6585 | 4020 | 0.61 | 1168 | 658 | 510 |
| Ca22chr1A *C. albicans* | 3188341 | 2348415 | 0.74 | 10927894 | 5463546 | 5464348 |
| Ca22chr1B *C. albicans* | 3188396 | 2260938 | 0.71 | 7163417 | 3581495 | 3581922 |
| Ca22chr2A *C. albicans* | 2231883 | 1686975 | 0.76 | 8851244 | 4422808 | 4428436 |
| Ca22chr2B *C. albicans* | 2231750 | 1630226 | 0.73 | 3421078 | 1709648 | 1711430 |
| Ca22chr3A *C. albicans* | 1799298 | 655796 | 0.36 | 2353987 | 1178205 | 1175782 |
| Ca22chr3B *C. albicans* | 1799271 | 639253 | 0.36 | 2568748 | 1284672 | 1284076 |
| Ca22chr4A *C. albicans* | 1603259 | 1230819 | 0.77 | 5433339 | 2717044 | 2716295 |
| Ca22chr4B *C. albicans* | 1603311 | 1227872 | 0.77 | 4929504 | 2463822 | 2465682 |
| Ca22chr5A *C. albicans* | 1190869 | 847851 | 0.71 | 3843038 | 1923191 | 1919847 |
| Ca22chr5B *C. albicans* | 1190991 | 847233 | 0.71 | 4512875 | 2255766 | 2257109 |
| Ca22chr6A *C. albicans* | 1033292 | 904473 | 0.88 | 5232771 | 2618032 | 2614739 |
| Ca22chr6B *C. albicans* | 1033212 | 902158 | 0.87 | 3665903 | 1833785 | 1832118 |
| Ca22chr7A *C. albicans* | 949580 | 299523 | 0.32 | 1174493 | 588246 | 586247 |
| Ca22chr7B *C. albicans* | 949611 | 293542 | 0.31 | 934591 | 466838 | 467753 |
| Ca22_Mitochondrion_ *C. albicans* | 40420 | 40126 | 0.99 | 9753599 | 4850600 | 4902999 |
| Ca22chrRA_ *C. albicans* | 2286237 | 1280174 | 0.56 | 6547437 | 3272552 | 3274885 |
| Ca22chrRB_ *C. albicans* | 2285697 | 1269869 | 0.56 | 4031665 | 2013982 | 2017683 |

**Table 3.** Mapping paired reads and coverage from raw sequencing data published by Muzzey et al. 2013 (Muzzey *et al.* 2013) to *C. albicans* (assembly 22, SC5314) and *S. epidermidis* ATCC 12228 (Zhang *et al.* 2003) reference sequences. Reads with non-specific matches were ignored and not included in the final mapping. The different plasmid samples of pSE-12228 correspond to the different GenBank sequences NC005008.1, NC005007.1, NC005006.1, NC005005.1, NC005004.1, NC005003.1 from top to bottom respectively.

| Reference Sequence | Reads in aligned pairs | Reads in broken pairs: wrong distance or mate inverted | Reads in broken pairs: mate on other contig | Reads in broken pairs: mate not mapped | Maximum coverage | Average coverage |
| --- | --- | --- | --- | --- | --- | --- |
| *S. epidermidis* chromosome | 40 | 5728 | 36048 | 233496 | 4925 | 1.95 |
| *S. epidermidis* plasmid pSE-12228-01 | 0 | 0 | 57 | 893 | 551 | 4.25 |
| *S. epidermidis* plasmid pSE-12228-02 | 0 | 0 | 99 | 398 | 69 | 1.96 |
| *S. epidermidis* plasmid pSE-12228-02 | 0 | 0 | 85 | 477 | 57 | 1.28 |
| *S. epidermidis* plasmid pSE-12228-02 | 0 | 0 | 480 | 2407 | 264 | 3.01 |
| *S. epidermidis* plasmid pSE-12228-02 | 0 | 0 | 414 | 2524 | 460 | 2.11 |
| *S. epidermidis* plasmid pSE-12228-02 | 0 | 0 | 121 | 1047 | 395 | 3.08 |
| Ca22chr1A *C. albicans* | 10558038 | 38800 | 40227 | 290829 | 16596 | 122.35 |
| Ca22chr1B *C. albicans* | 6950166 | 18398 | 31240 | 163613 | 2574 | 80.37 |
| Ca22chr2A *C. albicans* | 8601366 | 26490 | 26260 | 197128 | 8136 | 141.90 |
| Ca22chr2B *C. albicans* | 3280202 | 8872 | 21844 | 110160 | 2703 | 54.60 |
| Ca22chr3A *C. albicans* | 2259776 | 7888 | 11388 | 74935 | 3790 | 46.71 |
| Ca22chr3B *C. albicans* | 2486106 | 7176 | 10510 | 64956 | 3167 | 51.02 |
| Ca22chr4A *C. albicans* | 5256964 | 17628 | 25550 | 133197 | 2226 | 121.14 |
| Ca22chr4B *C. albicans* | 4764918 | 14510 | 24035 | 126041 | 3117 | 109.84 |
| Ca22chr5A *C. albicans* | 3716022 | 16510 | 15594 | 94912 | 5498 | 115.28 |
| Ca22chr5B *C. albicans* | 4384650 | 14882 | 15318 | 98025 | 1401 | 135.55 |
| Ca22chr6A *C. albicans* | 5033400 | 22224 | 23663 | 153484 | 4026 | 181.07 |
| Ca22chr6B *C. albicans* | 3489718 | 13638 | 21323 | 141224 | 27132 | 126.25 |
| Ca22chr7A *C. albicans* | 1121262 | 2922 | 7503 | 42806 | 5329 | 44.25 |
| Ca22chr7B *C. albicans* | 865670 | 3908 | 8976 | 56037 | 20173 | 35.17 |
| Ca22_Mitochondrion_ *C. albicans* | 9246170 | 304740 | 21989 | 180700 | 28525 | 8605.69 |
| Ca22chrRA_ *C. albicans* | 6340924 | 19000 | 28173 | 159340 | 21661 | 102.34 |
| Ca22chrRB_ *C. albicans* | 3863736 | 16226 | 24557 | 127146 | 2536 | 62.94 |

### Resistance to antifungal agents acquired in hypoxia condition in polymicrobial infections

Polymicrobial *C. albicans* cultures (Table 1) were tested for Minimal Inhibitory Concentrations (MIC). The results are summarized in Table 4. Strains SC5314 and L757 were pretreated with ampicillin to check whether the presence of the bacteria in cultures could influence the antifungal susceptibility profile.

**Table 4.** *In vitro* antifungal susceptibilities of *C. albicans* strains in normoxia and hypoxia at 37°C. Interval: concentrations (μg/ml) with growth. MIC_50_: Minimal concentration capable of 50% growth inhibition. MIC_90_: Minimal concentration capable of 90% growth inhibition. Classification according to the rules of the Clinical and laboratory Standards Institute (CLSI-M27-A3): Voriconazole (VRC) (S ≤ 1 μg/ml; SDD, 2 μg/ml; R ≥ 4 μg/ml) and Caspofugin (CAS) (S ≤ 0,25 μg/ml; I, 0,5 μg/ml; R ≥ 1 μg/ml). ^∗^na: not available.

| Conditions/Strains | MIC (µg/ml) |  |  |  |  |  |
| --- | --- | --- | --- | --- | --- | --- |
|  | Voriconazole (VRC) |  |  | Caspofugin (CAS) |  |  |
|  | Interval | MIC<sub>50</sub> | MIC<sub>90</sub> | Interval | MIC<sub>50</sub> | MIC<sub>90</sub> |
| **Hypoxia** |  |  |  |  |  |  |
| SC5314 | 0.00 | – | – | 0.01-0.25 | 0.13 | 0.25 |
| SC5314 (Ampicillin treated) | 0.03-16.00 | – | 16.00 | 0.01-0.25 | 0.13 | 0.25 |
| L757 | 0.00 | – | – | 0.01-0.50 | 0.25 | 0.50 |
| 9117 | 0.00 | – | – | 0.01-0.50 | 0.50 | 0.50 |
| ATCC 90029 | 0.00 | – | – | 0.01-1.00 | 0.50 | 1.00 |
| ATCC 24433 | 0.00 | – | – | 0.01-0.50 | 0.25 | 0.50 |
| **Normoxia** |  |  |  |  |  |  |
| SC5314 | 0.03-16.00 | – | 16.00 | 0.01-0.13 | 0.06 | 0.13 |
| SC5314 (Ampicillin treated) | 0.03-16.00 | – | 16.00 | 0.01-0.25 | 0.13 | 0.25 |
| L757 | 0.03-16.00 | – | 16.00 | 0.01-0.50 | 0.25 | 0.50 |
| 9117 | 0.03-16.00 | 16.00 | 16.00 | 0.01-0.50 | 0.50 | 0.50 |
| ATCC 90029 | 0.00 | – | – | 0.01-0.50 | 0.50 | 0.50 |
| ATCC 24433 | 0.03-16.00 | – | 16.00 | 0.01-0.13 | 0.06 | 0.13 |

We have observed that the *in vitro* evolution of SC5314 strain does not appear to interfere with the antifungal susceptibility profile when cells are grown in the absence of bacteria (data not shown). However, the presence of bacteria might affect the MIC90, in particular a drastic change in the susceptibility profile to Voriconazole (VRC) under hypoxia conditions, from resistant to sensitive (Table 4). The L757 strain also modified its susceptibility profile under hypoxia and in both absence and presence of bacteria, cells grown under hypoxia became sensitive to VRC. Although the other strains were not treated with ampicillin, they also presented the same profile (sensitive) under hypoxia, except for ATCC 90029 that seemed resistant both in normoxia and hypoxia. The antifungal Caspofungin (CAS), showed different inhibitory concentrations, however not enough to change the classification from resistant to sensitive. All strains were sensitive to CAS, except ATCC 90029, both in hypoxia and normoxia.

## DISCUSSION

*C. albicans* are commensal fungi and opportunistic pathogens capable of colonizing and/or infecting different niches of the human body. In addition to adapting to host conditions, such as temperature and low oxygen tension, these fungi interact with other microorganisms in the microbiota and can bind to them in polymicrobial infections that are commonly more severe than common candidemias and mortality rates up to twice as large (Pammi et al., 2013; Peleg et al., 2010).

In this work we have identified bacteria in single-colony derived *C. albicans* cultures. These cultures were started from an isolated colony of *C. albicans* SC5314, in plaques where no visual signs of bacterial contamination were detected. We were able to verify that even from an isolated colony it is possible to identify bacterial DNA by amplification of the 16S rRNA gene (Figure 1).

We observed that culture media, incubation period and temperature frequently used for *C. albicans* growth do not reveal bacterial growth, either by plating (Figure 2), by light microscopy or by 16S rRNA amplification (Figure 3). More importantly, in usual culture media, such as YPD and *CHROMagar*, although bacteria were not identified by plating or 16S rRNA amplification, it was possible to observe, by Gram staining, adhered bacterial cells especially in *C. albicans* hyphae (Figure 2). In addition, 16S rRNA gene amplification can easily lead to false-positive and false-negative results if not carried out with DNA-free high efficiency DNA polymerase with cycling conditions strictly tested and standardized for each negative control. For example, the increase in number of cycles can lead to amplification of a negative sample (without bacterial DNA added). These false-positives are due to contaminants in PCR reagents, including DNA polymerase enzymes (Carroll et al., 1999; Champlot et al., 2010; Heininger et al., 2003). The growth medium and incubation period were determinant for the appearance/identification of bacteria in our samples.

An *in vitro* evolution model was developed by our research group (Bartelli et al., 2017) and consists in serial culturing of *C. albicans* yeasts (strain SC5314) for prolonged periods under hypoxia, YPG medium (non-fermentative) at 37°C, (GTH strains). Results from this experiment showed a gradual increase in bacterial cells in aging cultures (Figure 4). These cultures were initiated by a single yeast colony from YPD plates incubated for 48h at 28°C. Among five *C. albicans* strains used in this study (Table 1) in different media, inoculation period and temperature (Figure 3), L757 was the only negative for bacterial 16S rRNA gene amplification during the 8-day incubation period, even in bacterial rich medium, such as LB (Figure 3). However, after 12 weeks of continuous growth in L757 GTH (data not shown), 16S rRNA amplification was positive. Sequencing of 16SrRNA gene revealed different bacterial species (Table 1). Strains L757 and ATCC 24433 showed *Propionibacterium acnes*, a slow growing anaerobic gram-positive bacteria. Recently *P. acnes* was recognized as an important opportunistic pathogen involved in invasive infections especially those associated with implants. This observation was possible due to improvement of isolation techniques, broad-range 16S rRNA sequencing and prolonged cultivation periods (up to 14 days) for periprosthetic biopsy specimens (Achermann et al., 2014; Portillo et al., 2013). The 16S rRNA sequencing of others strains (Table 1) revealed different species of genus *Staphylococcus*. Because the discrimination of this genera by 16S rRNA sequencing is controversial, sequencing of the *Tuf* gene was performed (Heikens et al., 2005) and readily revealed *S. epidermidis* in SC5324 cultures. The bacteria identified in the five *C. albicans* cultures tested (Table 1), as well as other bacteria of genera *Staphylococcus, Streptococcus, Pseudomonas* and *Corynebacterium* spp. are part of the microbiota in healthy individuals and also considered opportunistic pathogens. These species are especially involved in implant-mediated infections, forming mixed species biofilms that are associated with greater antimicrobial resistance and virulence (Achermann et al., 2014; Pierce et al., 2008; Pujol et al., 1993; Schlecht et al., 2015). Polymicrobial bloodstream infections are commonly associated with coagulase-negative Staphylococci (CNS), commonly *S. epidermidis*, and *Candida* species (Karlowicz et al., 2002; Pammi et al., 2013; Sutter et al., 2008). Bloodstream infections by *C. albicans* are associated with tissue invasion and staphylococcal infections are believed to be facilitated by these bacteria-hyphae associations. *S. epidermidis* can strongly adhere to *C. albicans* hyphae (~5nN), and also to its yeast forms (Beaussart et al., 2014; Pammi et al., 2013; Schlecht et al., 2015), which explains, at least in part, the frequent co-isolation of these two species and several results in this study.

**Figure 4.**
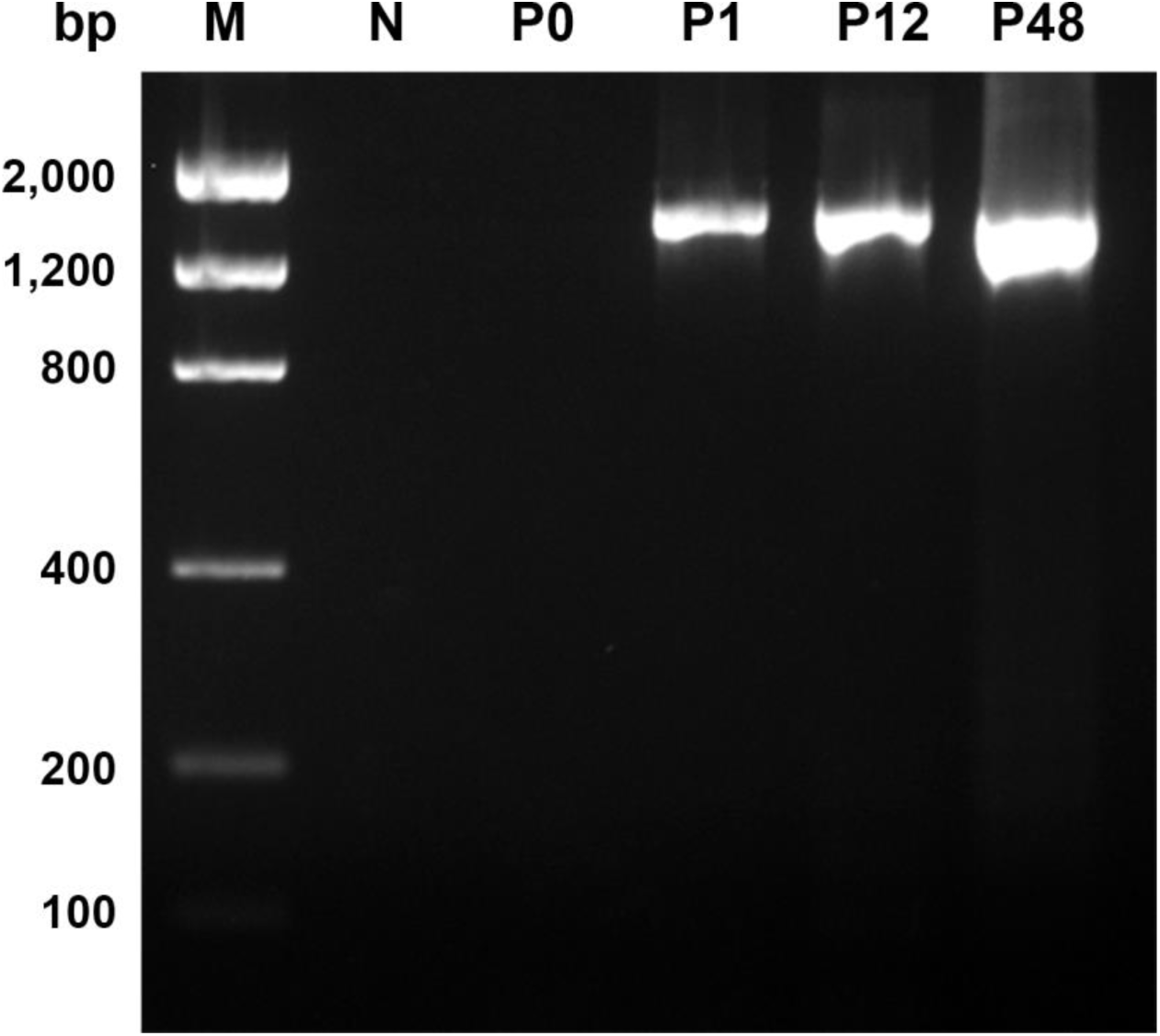
Amplification of the bacterial 16S rRNA gene in *C. albicans* SC5314 cultures during continuous growth (*in vitro* evolution experiments). Amplifications were carried out with an initial 200ng of total DNA and it was possible to observe an increase in the yield of 16S rRNA gene amplification along the continuous growth (weeks). **M**: Low mass ladder. **N**: negative control (no DNA added to the PCR reaction). **P0**: Cells grown in YPD for 48h at 28°C with agitation. **P1, P12** and **P48:** Cells grown continuously in YPG medium, 37°C in hypoxic environment (pO_2_ 5%) for 1, 12 and 48 weeks, respectively. Amplicons were visualized on 1% w/v agarose gel stained with ethidium bromide.

Consistent with another study (Klotz et al., 2007), we believe that *C. albicans* isolates can be associated with more than one bacterial species and/or genus. As shown by scanning electron microscopy (SEM), two different bacterial morphologies, coccus and bacillus can clearly be observed (Figure 5).

**Figure 5.**
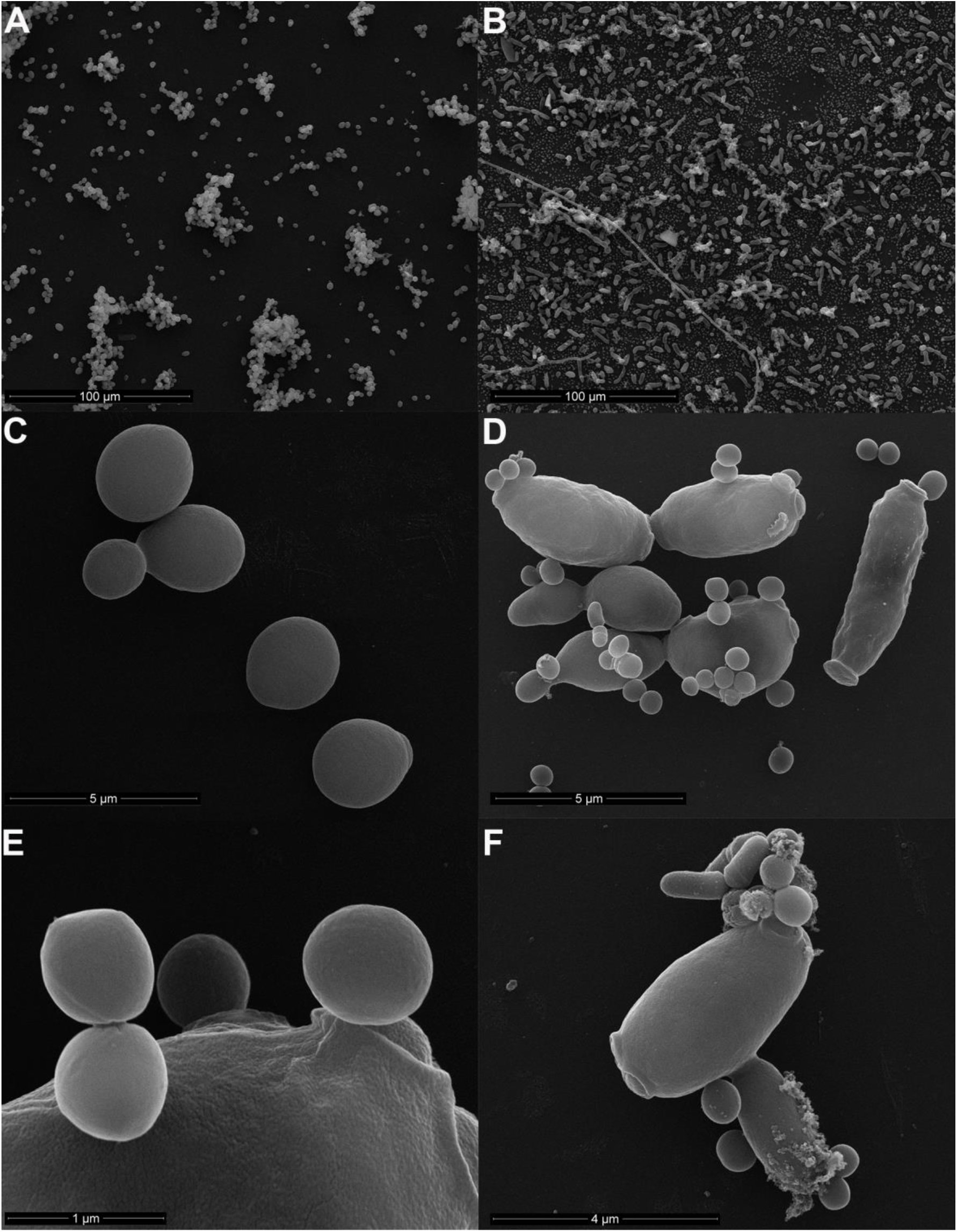
Bacteria adhered to *C. albicans* SC5314 visualized by scanning electron microscopy**. A** and **C**: Cultures were grown for 48h in YPD medium at 28°C under shacking. **B, D-F**: Cultures were grown at 37°C for 12 weeks in hypoxia and YPG medium. Scale bars are depicted in the pictures. **D** and **E**: Coccus adhered to the yeasts surface. **F**: Presence of more than one bacterial morphology, bacilli and coccus adhered to the yeast surface.

The fact that we identified bacterial reads in sequencing data from yeast cells, kept and manipulated in another laboratory, indicates that the yeast-bacteria co-isolation from single colonies might be underestimated and undetected when the appropriate medium and growth conditions are not used. In addition, our analysis of raw sequencing data from another lab (Tables 2 and 3) exclude the hypothesis that our *C. albicans* cultures were accidentally contaminated with *S. epidermidis* in our laboratory only. The same bacterial species, *S. epidermidis*, was identified both by the SC5314 strain present in our laboratory and in the laboratories involved in culturing the samples eventually used in the SC5314 genome sequencing (Muzzey et al., 2013). It is possible however that a single contamination event occurred in the very first isolation of SC5314 (in 1984) and therefore any cultures derived from the original SC5314 isolate would contain *S. epidermidis*.

It is estimated that 27-56% of candidemias are polymicrobial (Harriott and Noverr, 2009) and these are usually caused by CNS, most commonly *S. epidermidis* (Kloos and Bannerman, 1994; Pammi et al., 2013), which was found associated with two out of five strains tested here. We hypothesize that these bacterial-Candida associations are not more frequently observed because the majority of laboratories usually cultivate *Candida* for short periods of time and in bacteria inhibiting media. *C. albicans* isolation in clinical laboratories is performed for 24h to 72h, usually under yeast selective media, such as Sabouraud or *CHROMagar*^™^, previously shown to inhibit bacterial growth (Odds and Bernaerts, 1994; Sandven and Lassen, 1999; Vijaya et al., 2011). In addition, research laboratories, with few exceptions, cultivate yeasts for very short periods, usually overnight, during experimentation.

Polymicrobial infections caused by *C. albicans* and bacteria commonly involve the formation of biofilms, with adhesion of microorganisms to a substrate. *Staphylococcus* spp., in particular *S. aureus* and *S. epidermidis*, strongly interact with *C. albicans*, forming mixed biofilms and acquiring antimicrobial resistance (Pammi et al., 2013; Peters and Noverr, 2013; Schlecht et al., 2015).

To verify if the physical conditions that mimic the host could influence *C. albicans* drug-resistance, the MICs of polymicrobial *C. albicans* cultures (Table 1) were tested. The results summarized in Table 4 show that the presence of bacteria affects the MICs, in particular a drastic change in the susceptibility profile to VRC under hypoxia. In addition, the antifungal CAS showed different inhibitory concentrations, but not enough to change the standardized classification from resistant to sensitive. All strains were classified as CAS sensitive, except for ATCC 90029, both in hypoxia and normoxia. However, we could observe an increase in CAS concentrations in hypoxia for ATCC 90029.

Resistance to antifungal is a major problem, resulting in a drastic increase in the incidence of opportunistic and systemic fungal infections, especially in immunocompromised individuals. There is a limited range of antifungal agents to treat infections caused by *Candida* spp., including polyenes, flucytosine, azoles and echinocandins, and a variety of widely accepted techniques as reference methods, especially CLSI-M27-A3 (Warn et al., 2004 CLSI 2008). However, most techniques do not take into account for growth conditions in the host. In the human body the oxygen tension is around 2-9%, the temperature is 37°C and *C. albicans* is in constant presence of other microbiota microorganisms. Two studies suggest, however, that there are no major differences between *in vitro* and *in vivo* resistance for some antifungal agents (Grahl et al., 2012; Warn et al., 2004). Nevertheless, in face of data here presented, testing for drug resistance might benefit by culturing microorganisms in conditions that are as close as possible to the conditions inside the host.

Azoles are part of an important class of antifungal agents commonly used as a gold standard against infections caused by *Candida* spp. However, the increase in its use, coupled with the fact that they are fungistatic drugs, led to the emergence of resistant isolates of *Candida* spp. (Vandeputte et al., 2009). Although the vast majority of studies suggest that hypoxia promotes increased drug resistance, especially to azoles (Dumitru et al., 2007), we observed that *C. albicans* strains under hypoxia presented a sensitivity profile to VRC. Several authors have already reported the hypersensitivity profile to azoles. In *C. albicans*, deletion of *CaUPC2*, a transcriptional factor, decreases ergosterol, as well as an impairment in expression regulation of genes involved in ergosterol biosynthesis. Damage to *C. albicans* cell wall might also favor increased susceptibility as well as deficiency in the respiratory chain (Silver et al., 2004; Sun et al., 2013; Vandeputte et al., 2009).

Although the susceptibility profile to VRC changed from resistant to sensitive under hypoxia, the SC5314 strain treatment with ampicillin did not present this phenotype (Table 4). We believe that the presence of *Staphylococcus* CNS, more specifically *S. epidermidis*, might have interfered. The association of *C. albicans* infections with *S. epidermidis* is known to difficult the treatment by azoles. We do not believe that the change in the susceptibility profile is due to the strain SC5314ampicillin treatment because strain L757 when treated in the same manner, had a hypoxia sensitive profile both in the absence (ampicillin treated cells) and in the presence of bacteria. In addition, there is no data showing that ampicillin interferes with *C. albicans* growth.

Sensitivity to VRC under hypoxia by SC5314 may be related to intrinsic characteristics of the yeast itself and to the interaction with *S. epidermidis*. The main quorum sensing molecule produced by of C. *albicans* is farnesol, which is able to mediate intra and interspecies interactions (De Sordi and Mühlschlegel, 2009). Farnesol is generated endogenously by farnesyl pyrophosphate (FPP), affects the expression of genes involved in the cellular response to stress, thermal shock, antimicrobial resistance, metabolism, cell wall and cell cycle. FPP is a precursor of ergosterol synthesis and sterol biosynthesis, thus the azoles interfere in the intra and extracellular levels of farnesol (Shareck and Belhumeur, 2011; Yu et al., 2012). In this way we believe that the sensitivity conferred only in the presence of bacteria for SC5314 can be derived from farnesol or other quorum sensing molecules, and this result can be corroborated by Yu et al. (2012) (Yu et al., 2012) who demonstrate a correlation between farnesol and ergosterol biosynthesis acting as a signaling molecule in regulating the expression of ERG genes and efflux pumps, resulting in an inhibition of fluconazole resistance in C. *albicans*. However, mixed biofilms of *C. albicans* and *S. epidermidis* are known to have a decreased antifungal susceptibility through slime produced by *S. epidermidis* (Adam et al., 2002). Therefore, hypoxia is essential to determine the change in the susceptibility profile to antifungal agents and should be taken into account in clinical treatment and routine susceptibility tests. Although further studies are needed to comprehend the bacterial-fungal interaction, determining whether cultures of *C. albicans* are monomicrobial or polymicrobial have a direct impact on the fungus survival and its virulence, especially concerning patient management and treatment options.

## MATERIALS AND METHODS

### *C. albicans* strains and growth conditions

We used five *C. albicans* strains. Strain SC5314, kindly provided by Dr A. Mitchell, Carnegie Mellon University, and strains L757, 9117, ATCC 24433 and 90029 kindly provided by A.L. Colombo, Departamento de Infectologia, Laboratório Especial de Micologia – Unifesp (Table 1). Strains were grown under YPD (1% yeast extract, 2% peptone, 2% dextrose and 2% agar when needed), YPG (1% yeast extract, 2% Bacto-peptone, 2% glycerol, and 2% agar when needed) and LB (1% yeast extract, 0.5% Bacto-tryptone, 1% NaC1 and 2% agar for plates) for 48 hours on agar plates, and for 48h or 8 days under liquid media. Cells were inoculated into 10 mL of fresh medium and samples were incubated at different temperatures (28°C or 37°C) and oxygen conditions, such as normóxia (agitation 150 rpm) or hypoxia (cells were maintained in hermetic closed jars with a hypoxia generator Microaerobac, Probac do Brasil - with an atmosphere of 5-15% O2). The strain SC5314 was also grown on blood agar plates (Probac do Brasil) and *CHROMagar* (Difco) for 48h at 37°C.

### Visual inspection of bacteria in *C. albicans* cultures

To verify if bacteria were present in the cultures grown for 48h or 8 days under different growth conditions, agar plates (YPD, LB, *CHROMagar* and blood agar) were inspected after 48h growth at 37°C or 28°C. Yeast colonies were also inspected by light microscopy with under 100X oil immersion objective (colonies were collected from different growth media for 48h and/or 8 days from one isolated colony) and Gram staining (Hucker, 1921).

### Scanning electron microscopy

Bacteria were detected in cultures of *C. albicans* SC5314 by Scanning Electron Microscopy (SEM). Cells were washed with saline solution and fixated in 2.5% glutaraldehyde and 0.1M sodium cacodylate buffer pH 7.2 for 2h in room temperature. Samples were washed with 0.1M sodium cacodylate (2x for 10 min, 1x overnight and 1x for 10 min). Cells were incubated in 1% osmium tetroxide and 0.1M sodium cacodylate for 1h and washed 3x for 10 min in the sodium cacodylate buffer. Cells were treated in 1% tannic acid for 30min, washed 3x for 5min in water. Cells were dehydrated in ethanol (2x 50% 10min, 2x 70% 10min, 2x 90% 10min and 3x 100% 10 min). Samples were dried and covered with gold by sputtering 20-30nm (©LEICA at SCD 500, Germany) before observation by Field Issue Quanta FEG 250.

### Detection and identification of bacteria by 16S rRNA gene amplification and sequencing

DNA extractions were carried out by two different protocols, one aiming to extract total yeast DNA (Wach et al., 1994) and the other modified for bacterial DNA extraction (Wilson, 2001). Sequences of primers used for amplification of 16S rRNA gene were extracted from (Sauer et al., 2005). The amplification reactions consisted of 25μl containing 12.5μl of 2X My Taq Master Mix (Bioline), 0.5 μM of each forward and reverse primer and 50ng of DNA with cycling conditions of 95°C for 1 min, 30 cycles of 95°C 15s, 58°C 15s, and 72°C 30s. For *Staphylococcus* spp. discrimination, the gene *tuf* was also amplified with a 20μl reaction volume containing 10μl of 2X Master Mix (Fermentas), 0.5 μM of each forward and reverse primer (Heikens et al., 2005) and 50ng of DNA with 95°C 5 min, 30 cycles of 95°C 1 min, 55°C 1 min, 72°C 1 min and final extension of 72°C for 10 min. The DNA used for amplification was derived from yeast and/or bacterial extraction protocols. Amplicons were sequenced on both strands with an ABI Prism 3130xl automated sequencer (Applied Biosystems). Reads were *de novo* assembled with Geneious v.6.1.5 (Biomatters Ltd.), and low quality regions in the 5’ and 3’ ends of the consensus sequence with eventual ambiguities and low quality base calling (phred score <20) were removed. To identify the bacterial species, the consensus sequences were subjected to a search on the Nucleotide BLAST tool (http://blast.ncbi.nlm.nih.gov) using the 16S ribosomal RNA Database (Bacteria and Archaea) optimized for GenBank Highly similar sequences (megablast). The criteria for species identification followed the recommendations described in Janda and Abbott (2007) (Janda and Abbott, 2007), with a minimum of 99%-99.5% sequence similarity between query and subject.

### Analysis of *C. albicans* SC5314 raw genome sequencing data

Sequencing reads were downloaded from SRA BioProject SRP022363 (Muzzey et al., 2013) and mapped simultaneously to the *C. albicans* SC5314 reference sequence (version A22-s05-m04-r02, in http://www.Candidagenome.org) and *S. epidermidis* ATCC 12228 (chromosome NC_004461.1, and plasmids NC_005008.1, NC_005007.1, NC_005006.1, NC_005005.1, NC_005004.1, NC_005003.1) with CLC Genomics Workbench v.7.5.1 (Qiagen) with default parameters except for ignoring and excluding reads with non-specific matches.

### Experimental evolution

For the experimental evolution, cells of strains SC5314 and L757 were grown on YPD plate (1% w/v yeast extract; 2% w/v peptone, 2% w/v dextrose, 2% w/v agar) and a single colony was used for overnight growth on YPD broth at 28°C, 150 rpm. 1x105 cells of this inoculums were added into tubes containing 10mL of fresh YPD (1% w/v yeast extract; 2% w/v peptone, 2% w/v dextrose) or YPG (1% w/v yeast extract; 2% w/v peptone, 2% w/v glycerol). Cells were cultivated continuously at presence of bacterial (Table 1) in different temperatures (28°C or 37°C) and oxygen availability conditions (normoxia or hypoxia) for 12 weeks. Once a week, 0.1% of cell cultures were transferred to fresh 10mL medium for created an evolution in vitro. To generate a hypoxic environment, cells were maintained in hermetic closed jars with a hypoxia generator (Microaerobac, Probac do Brasil) with an atmosphere of 5-15% O2, while cells under normoxia were kept under 150 rpm agitation with O2 levels close to 21% (Bartelli et al., 2017). The other strains 9117, ATCC 24433 and ATCC 90029 of C. *albicans*, grown on YPD broth at 28°C, 150 rpm for 48 hours, only to be performed to antifungal susceptibility testing.

### Ampicillin treatment

Strains SC5314 and L757 were first plated on YPD with 50mg/ml ampicillin, after 48h a single colony was selected and added in 10 ml of YPD medium with 20 mg/ml ampicillin. Cells were grown for one week at 28°C at 150 rpm, and during that week every two days 20 mg/ml ampicillin was added to ensure that there would be no more bacteria. Cultures were washed with sterile PBS 1X and incubated without ampicillin, under the appropriate conditions of experimental evolution. Absence of bacteria was confirmed by negative 16S rRNA amplification.

### In vitro antifungal susceptibility testing

Antifungal susceptibility testing was performed using the broth microdilution method according to the Clinical and Laboratory Standards Institute (CLSI M27-A3/S4) (CLSI, 2008, 2012). Antifungals tested were Voriconazole (VRC) and Caspofungin (CAS) (Sigma Chemical Corporation St. Louis, MO). Assays were incubated at 37°C in normoxia and hypoxia conditions. ATCC 24433 was used as quality control, following the CLSI M27-A3 guidelines. The Minimal Inhibitory Concentrations (MIC) of antifungals were determined by visual readings and defined as the lowest concentration capable of inhibiting 50% and 90% of cell growth. Antifungal concentrations tested were: 0.015-8.00 μg/ml (CAS) (S ≤ 0,25 μg/ml; I, 0,5 μg/ml; R ≥ 1 μg/ml) and 0.031-16.00 μg/ml (VRC) Voriconazole (S ≤ 1 μg/ml; SDD, 2 μg/ml; R ≥ 4 μg/ml). Assays were performed in triplicates in at least two independent experiments.

## ACKNOWLEDGMENTS

We thank Prof. Arnaldo Lopes Colombo, Departamento de Infectologia, Laboratório Especial de Micologia (LEMI) – Unifesp for providing *C. albicans* clinical isolates. We also thank, Centro de Microscopia Eletrônica (CEME) – Unifesp by performing the SEM.

## AUTHOR CONTRIBUTIONS

D.C.F.B. planned and performed experiments, analyzed data, discussed the results and implications and wrote the manuscript. T.F.B.: planned experiments, performed sequencing and bioinformatics analysis, discussed the results and implications, and edited the manuscript. C.R.R. helped technically with MICs. M.R.S.B. planned and supervised experiments, wrote and edited the manuscript.

## REFERENCES

Achermann, Y., Goldstein, E. J. C., Coenye, T., and Shirtliff, M. E. (2014). Propionibacterium acnes: from commensal to opportunistic biofilm-associated implant pathogen. Clin. Microbiol. Rev. 27, 419–440. doi:10.1128/CMR.00092-13.

Adam, B., Baillie, G. S., and Douglas, L. J. (2002). Mixed species biofilms of Candida albicans and Staphylococcus epidermidis. J. Med. Microbiol. 51, 344–349. doi:10.1099/0022-1317-51-4-344.

Arumugam, M., Raes, J., Pelletier, E., Le Paslier, D., Yamada, T., Mende, D. R., et al. (2011). Enterotypes of the human gut microbiome. Nature 473, 174–180. doi:10.1038/nature09944.

Bartelli, T. F., Bruno, D.C., Lichtenstein, F., and Briones, M.R.S. (2017). Experimental evolution of Candida albicans under hypoxia and heat shock reveals nuclear genome variants and mitochondrial methylome alterations. Biorxiv, doi: 10.1101/167338.

Beaussart, A., El-Kirat-Chatel, S., Sullan, R. M. A., Alsteens, D., Herman, P., Derclaye, S., et al. (2014). Quantifying the forces guiding microbial cell adhesion using single-cell force spectroscopy. Nat. Protoc. 9, 1049–1055. doi:10.1038/nprot.2014.066.

Bergman, A., and Casadevall, A. (2010). Mammalian Endothermy Optimally Restricts Fungi and Metabolic Costs. mBio 1, e00212-10. doi:10.1128/mBio.00212-10.

Brock, M. (2009). Fungal metabolism in host niches. Curr. Opin. Microbiol. 12, 371–376. doi:10.1016/j.mib.2009.05.004.

Brown, G. D., Denning, D. W., Gow, N. A. R., Levitz, S. M., Netea, M. G., and White, T. C. (2012). Hidden Killers: Human Fungal Infections. Sci. Transl. Med. 4, 165rv13-165rv13. doi:10.1126/scitranslmed.3004404.

Carroll, N. M., Adamson, P., and Okhravi, N. (1999). Elimination of bacterial DNA from Taq DNA polymerases by restriction endonuclease digestion. J. Clin. Microbiol. 37, 3402–3404.

Champlot, S., Berthelot, C., Pruvost, M., Bennett, E. A., Grange, T., and Geigl, E.-M. (2010). An efficient multistrategy DNA decontamination procedure of PCR reagents for hypersensitive PCR applications. PloS One 5. doi:10.1371/journal.pone.0013042.

CLSI (2008). M27-A3Reference Method for Broth Dilution Antifungal Susceptibility Testing of Yeasts, 3rd Edition.

CLSI (2012). M27-S4Reference Method for Broth Dilution Antifungal Susceptibility Testing of Yeasts; Fourth Informational Supplement. 4th ed.

De Sordi, L., and Mühlschlegel, F. A. (2009). Quorum sensing and fungal-bacterial interactions in Candida albicans: a communicative network regulating microbial coexistence and virulence. FEMS Yeast Res. 9, 990–999.

Douglas, L. J. (2002). Medical importance of biofilms in Candida infections. Rev. Iberoam. Micol. 19, 139–143.

Dumitru, R., Navarathna, D. H. M. L. P., Semighini, C. P., Elowsky, C. G., Dumitru, R. V., Dignard, D., et al. (2007). In vivo and in vitro anaerobic mating in Candida albicans. Eukaryot. Cell 6, 465–472. doi:10.1128/EC.00316-06.

Espinel-Ingroff, A., Kish, C. W., Kerkering, T. M., Fromtling, R. A., Bartizal, K., Galgiani, J. N., et al. (1992). Collaborative comparison of broth macrodilution and microdilution antifungal susceptibility tests. J. Clin. Microbiol. 30, 3138–3145.

Gillum, A. M., Tsay, E. Y., and Kirsch, D. R. (1984). Isolation of the Candida albicans gene for orotidine-5’-phosphate decarboxylase by complementation of S. cerevisiae ura3 and E. coli pyrF mutations. Mol. Gen. Genet. MGG 198, 179–182.

Grahl, N., Shepardson, K. M., Chung, D., and Cramer, R. A. (2012). Hypoxia and fungal pathogenesis: to air or not to air? Eukaryot. Cell 11, 560–570. doi:10.1128/EC.00031-12.

Harriott, M. M., and Noverr, M. C. (2009). Candida albicans and Staphylococcus aureus form polymicrobial biofilms: effects on antimicrobial resistance. Antimicrob. Agents Chemother. 53, 3914–3922. doi:10.1128/AAC.00657-09.

Heikens, E., Fleer, A., Paauw, A., Florijn, A., and Fluit, A. C. (2005). Comparison of genotypic and phenotypic methods for species-level identification of clinical isolates of coagulase-negative staphylococci. J. Clin. Microbiol. 43, 2286–2290. doi:10.1128/JCM.43.5.2286-2290.2005.

Heininger, A., Binder, M., Ellinger, A., Botzenhart, K., Unertl, K., and Döring, G. (2003). DNase pretreatment of master mix reagents improves the validity of universal 16S rRNA gene PCR results. J. Clin. Microbiol. 41, 1763–1765.

Hucker, G. J. (1921). A New Modification and Application of the Gram Stain. J. Bacteriol. 6, 395–397.

Janda, J. M., and Abbott, S. L. (2007). 16S rRNA gene sequencing for bacterial identification in the diagnostic laboratory: pluses, perils, and pitfalls. J. Clin. Microbiol. 45, 2761–2764. doi:10.1128/JCM.01228-07.

Karlowicz, M. G., Furigay, P. J., Croitoru, D. P., and Buescher, E. S. (2002). Central venous catheter removal versus in situ treatment in neonates with coagulase-negative staphylococcal bacteremia. Pediatr. Infect. Dis. J. 21, 22–27.

Kloos, W. E., and Bannerman, T. L. (1994). Update on clinical significance of coagulase-negative staphylococci. Clin. Microbiol. Rev. 7, 117–140.

Klotz, S. A., Gaur, N. K., De Armond, R., Sheppard, D., Khardori, N., Edwards, J. E., et al. (2007). Candida albicans Als proteins mediate aggregation with bacteria and yeasts. Med. Mycol. 45, 363–370. doi:10.1080/13693780701299333.

Mayer, F. L., Wilson, D., and Hube, B. (2013). Candida albicans pathogenicity mechanisms. Virulence 4, 119–128. doi:10.4161/viru.22913.

Muzzey, D., Schwartz, K., Weissman, J. S., and Sherlock, G. (2013). Assembly of a phased diploid Candida albicans genome facilitates allele-specific measurements and provides a simple model for repeat and indel structure. Genome Biol. 14, R97. doi:10.1186/gb-2013-14-9-r97.

Odds, F. C., and Bernaerts, R. (1994). CHROMagar Candida, a new differential isolation medium for presumptive identification of clinically important Candida species. J. Clin. Microbiol. 32, 1923–1929.

Ovchinnikova, E. S., Krom, B. P., Busscher, H. J., and van der Mei, H. C. (2012). Evaluation of adhesion forces of Staphylococcus aureus along the length of Candida albicans hyphae. BMC Microbiol. 12, 281. doi:10.1186/1471-2180-12-281.

Pammi, M., Liang, R., Hicks, J., Mistretta, T.-A., and Versalovic, J. (2013). Biofilm extracellular DNA enhances mixed species biofilms of Staphylococcus epidermidis and Candida albicans. BMC Microbiol. 13, 257. doi:10.1186/1471-2180-13-257.

Peleg, A. Y., Hogan, D. A., and Mylonakis, E. (2010). Medically important bacterial-fungal interactions. Nat. Rev. Microbiol. 8, 340–349. doi:10.1038/nrmicro2313.

Peters, B. M., and Noverr, M. C. (2013). Candida albicans-Staphylococcus aureus polymicrobial peritonitis modulates host innate immunity. Infect. Immun. 81, 2178-2189. doi:10.1128/IAI.00265-13.

Pfaller, M. A., Messer, S. A., and Hollis, R. J. (1994). Strain delineation and antifungal susceptibilities of epidemiologically related and unrelated isolates of Candida lusitaniae. Diagn. Microbiol. Infect. Dis. 20, 127–133.

Pierce, C. G., Uppuluri, P., Tristan, A. R., Wormley, F. L., Mowat, E., Ramage, G., et al. (2008). A simple and reproducible 96-well plate-based method for the formation of fungal biofilms and its application to antifungal susceptibility testing. Nat. Protoc. 3, 1494–1500. doi:10.1038/nport.2008.141.

Pierce, G. E. (2005). Pseudomonas aeruginosa, Candida albicans, and device-related nosocomial infections: implications, trends, and potential approaches for control. J. Ind. Microbiol. Biotechnol. 32, 309–318. doi:10.1007/s10295-005-0225-2.

Portillo, M. E., Corvec, S., Borens, O., and Trampuz, A. (2013). Propionibacterium acnes: an underestimated pathogen in implant-associated infections. BioMed Res. Int. 2013, 804391. doi:10.1155/2013/804391.

Pujol, C., Reynes, J., Renaud, F., Raymond, M., Tibayrenc, M., Ayala, F. J., et al. (1993). The yeast Candida albicans has a clonal mode of reproduction in a population of infected human immunodeficiency virus-positive patients. Proc. Natl. Acad. Sci. U. S. A. 90, 9456–9459.

Rodaki, A., Bohovych, I. M., Enjalbert, B., Young, T., Odds, F. C., Gow, N. A. R., et al. (2009). Glucose promotes stress resistance in the fungal pathogen Candida albicans. Mol. Biol. Cell 20, 4845–4855. doi:10.1091/mbc.E09-01-0002.

Sandven, P., and Lassen, J. (1999). Importance of selective media for recovery of yeasts from clinical specimens. J. Clin. Microbiol. 37, 3731–3732.

Sauer, P., Gallo, J., Kesselová, M., Kolár, M., and Koukalová, D. (2005). Universal primers for detection of common bacterial pathogens causing prosthetic joint infection. Biomed. Pap. Med. Fac. Univ. Palacky Olomouc Czechoslov. 149, 285–288.

Schlecht, L. M., Peters, B. M., Krom, B. P., Freiberg, J. A., Hänsch, G. M., Filler, S. G., et al. (2015). Systemic Staphylococcus aureus infection mediated by Candida albicans hyphal invasion of mucosal tissue. Microbiol. Read. Engl. 161, 168–181. doi:10.1099/mic.0.083485-0.

Setiadi, E. R., Doedt, T., Cottier, F., Noffz, C., and Ernst, J. F. (2006). Transcriptional response of Candida albicans to hypoxia: linkage of oxygen sensing and Efg1p-regulatory networks. J. Mol. Biol. 361, 399–411. doi:10.1016/j.jmb.2006.06.040.

Shareck, J., and Belhumeur, P. (2011). Modulation of morphogenesis in Candida albicans by various small molecules. Eukaryot. Cell 10, 1004–1012. doi:10.1128/EC.05030-11.

Silver, P. M., Oliver, B. G., and White, T. C. (2004). Role of Candida albicans transcription factor Upc2p in drug resistance and sterol metabolism. Eukaryot. Cell 3, 1391–1397. doi:10.1128/EC.3.6.1391-1397.2004.

Sims, C. R., Ostrosky-Zeichner, L., and Rex, J. H. (2005). Invasive candidiasis in immunocompromised hospitalized patients. Arch. Med. Res. 36, 660–671. doi:10.1016/j.arcmed.2005.05.015.

Sun, N., Fonzi, W., Chen, H., She, X., Zhang, L., Zhang, L., et al. (2013). Azole susceptibility and transcriptome profiling in Candida albicans mitochondrial electron transport chain complex I mutants. Antimicrob. Agents Chemother. 57, 532–542. doi:10.1128/AAC.01520-12.

Sutter, D., Stagliano, D., Braun, L., Williams, F., Arnold, J., Ottolini, M., et al. (2008). Polymicrobial bloodstream infection in pediatric patients: risk factors, microbiology, and antimicrobial management. Pediatr. Infect. Dis. J. 27, 400–405. doi:10.1097/INF.0b013e31816591be.

Vandeputte, P., Tronchin, G., Rocher, F., Renier, G., Bergès, T., Chabasse, D., et al. (2009). Hypersusceptibility to azole antifungals in a clinical isolate of Candida glabrata with reduced aerobic growth. Antimicrob. Agents Chemother. 53, 3034–3041. doi:10.1128/AAC.01384-08.

Vijaya, D., Harsha, T. R., and Nagaratnamma, T. (2011). Candida speciation using chrom agar. 5, 755–757.

Vilanova, M., and Correia, A. (2008). Host defense mechanisms in invasive candidiasis originating in the GI tract. Expert Rev. Anti Infect. Ther. 6, 441–445. doi:10.1586/14787210.6.4.441.

Wach, A., Brachat, A., Pöhlmann, R., and Philippsen, P. (1994). New heterologous modules for classical or PCR-based gene disruptions in Saccharomyces cerevisiae. Yeast Chichester Engl. 10, 1793–1808.

Warn, P. A., Sharp, A., Guinea, J., and Denning, D. W. (2004). Effect of hypoxic conditions on in vitro susceptibility testing of amphotericin B, itraconazole and micafungin against Aspergillus and Candida. J. Antimicrob. Chemother. 53, 743–749. doi:10.1093/jac/dkh153.

Wilson, K. (2001). Preparation of genomic DNA from bacteria. Curr. Protoc. Mol. Biol. Chapter 2, Unit 2.4. doi:10.1002/0471142727.mb0204s56.

Yu, L.-H., Wei, X., Ma, M., Chen, X.-J., and Xu, S.-B. (2012). Possible inhibitory molecular mechanism of farnesol on the development of fluconazole resistance in Candida albicans biofilm. Antimicrob. Agents Chemother. 56, 770–775. doi:10.1128/AAC.05290-11.

Zhang, Y.-Q., Ren, S.-X., Li, H.-L., Wang, Y.-X., Fu, G., Yang, J., et al. (2003). Genome-based analysis of virulence genes in a non-biofilm-forming Staphylococcus epidermidis strain (ATCC 12228). Mol. Microbiol. 49, 1577–1593.

